# SeqOthello: Query over RNA-seq experiments at scale

**DOI:** 10.1101/258772

**Authors:** Ye Yu, Jinpeng Liu, Xinan Liu, Yi Zhang, Eamonn Magner, Chen Qian, Jinze Liu

**Affiliations:** University of Kentucky; University of California Santa Cruz

## Abstract

We present SeqOthello, an ultra-fast and memory-efficient indexing structure to support arbitrary sequence query against large collections of RNA-seq experiments. SeqOthello requires only five minutes to conduct a global survey of 11,658 fusion events against 10,113 TCGA Pan-Cancer RNA-seq datasets on a standard computer with 19.1 GB memory space. The query recovers 92.7% of tier-1 fusions curated by TCGA Fusion Gene Database and further reveals 270 novel fusion occurrences, all of which present as tumor-specific. The entire index is only 76 GB, achieving a 700:1 compression ratio relative to the original sequencing data and making it extremely portable. This is the first sequence search index constructed on the scale of TCGA data. By providing a reference-free, alignment-free, and parameter-free sequence search system, SeqOthello will enable large-scale integrative studies using sequence-level data, an undertaking not previously practicable for many individual labs. SeqOthello is currently available at https://github.com/LiuBioinfo/SeqOthello.

## Introduction

Advances in the study of functional genomics over the past decade have produced a vast supply of RNA-seq datasets. As of December 2017, over 12 Petabytes of RNA-seq data were deposited in the Sequence Read Archive (SRA)^1^. Sequencing consortiums such as The Cancer Genome Atlas (TCGA)^2^ and the International Cancer Genomics Consortium (ICGC)^3^ have sequenced tens of thousands of tumor transcriptomes from a variety of cancer populations. Although these datasets have collectively redefined the landscape of cancer transcriptome, more features relevant from a clinical perspective remain to be discovered. However, data reanalysis requires extensive computational resources and bioinformatics support, making it exclusive to a few labs. The development of SeqOthello will enable many labs with limited resources to learn from sequencing-level data by supporting fast and memory-efficient query over large-scale RNA-seq datasets.

To date, sequence search options are limited. Most sequencing databases support meta-data searches^1,3,4^, which permit selection of experiments by tissue type, organism, experimental condition or sequencing protocol. From this refined list, experiments can be downloaded and analyzed individually^5^. Alternately, SRA-BLAST^6^ can retrieve short reads aligned to a query sequence, but only for a limited number of nucleotides per query. Finally, the Bioinformatics community has lately established databases storing ready-to-analyze results in areas such as gene expression^4,7,8^ and exon-exon junctions^9^. However, these databases are subject to frequent updates as Bioinformatics algorithms improve and reference genomes are refined, nor can they support the query of novel sequences that are absent from existing annotation or undetectable by current bioinformatics tools.

Recently, Sequence Bloom Tree (SBT)^10^ and its descendants^11,12^ were developed to query RNA-seq experiments for expressed transcripts, pioneering the field of large-scale sequence search in RNA-seq. SBT is designed as an experiment filter that returns the subset of experiments containing at least *θ* percent of *k*-mers from the query sequence. Built upon bloom filters^13,14^, SBT-based algorithms are generally memory efficient for small queries. Unfortunately, tuning the input parameter *θ* is time-consuming and produces inconsistent results for a single query, thereby hampering interpretability. Furthermore, extracting sequence-level information from the filtered experiments requires downloading and reanalyzing of raw sequencing datasets, and thus does not sidestep traditional RNA-seq processing. There is also growing interest in methods for indexing large collections of genomic sequencing reads from different individuals. Bloom filter trie (BFT)^15^ was developed to store and compress a set of colored k-mers from a Pan-Genome of hundreds of samples. Additionally, the Burrows–Wheeler transform (BWT) and FM-index have been employed to build indexes on raw sequencing reads with applications in compressing 2705 whole genome sequencing samples from the 1000 Genomes Project^16,17^. Though retaining full-text information, these data structures are often associated with high memory cost and slow query speed as the entire index must be loaded to memory prior to query.

Here we present SeqOthello, a novel indexing structure that supports query of an arbitrary sequence against large collections of RNA-seq experiments. Large-batch query with SeqOthello is orders of magnitude faster than with SSBT, the improved version of SBT. A SeqOthello query may return near-exact *k*-mer information in individual experiments or *k*-mer hit ratios (*i.e.*, the fraction of *k*-mer hits in a query). We illustrate the utility and efficiency of SeqOthello by conducting a global survey of known gene fusions against 10,113 TCGA RNA-seq datasets. The survey confirms roughly 93% of known fusion events and reveals close to 300 novel fusion occurrences, all of which are tumor-specific. The entire survey was completed in under five minutes and required a computer with no more than 32 GB memory, which, to our knowledge, is a scale unachieved by previous methods.

## Results

### SeqOthello Data Structure

A sequencing experiment can be represented by a collection of *k*-mers, or length *k* subsequences of the original reads. *k*-mers are fundamental components of de Bruijn graphs and thus are essential for *de novo* assembly of the transcriptome^18–20^ in individual experiments. A *database* of sequencing experiments can therefore be represented as the *occurrence map* of individual *k*-mers, defined as the presence or absence of these k-mers in each experiment. The challenge is to efficiently store and query this information in scenarios with billions of *k*-mers across tens of thousands of experiments. We leverage novel algorithms in data compression and *k*-mer indexing to surmount this obstacle.

The prevalence of each *k*-mer varies dramatically, with plots often exhibiting a *U*- or *L*-shaped distribution (Supp Figure 1). *k*-mers located at the extremes of the spectrum tend to originate from experiment-specific transcripts, or to descend from housekeeping genes, and thus manifest in nearly all experiments. By contrast, *k*-mers near the center of the distribution may be tissue- or organism-specific. The prevalence of a *k*-mer directly determines the information content^21,22^, or the number of bits required to store its occurrence map. To this end, SeqOthello employs an information-content-aware data-compression scheme: an ensemble of compression techniques tailored to store the occurrence maps of *k*-mers from each region of the occurrence distribution, without hampering query efficiency (Figure. 1a. and Methods). SeqOthello relies on a novel, hierarchical indexing structure to facilitate fast retrieval of *k*-mer occurrence maps (Figure 1 a). The mappings between levels are supported by the *Othello*^23,24^ data structure (Methods), a minimal perfect hashing classifier that provides key-to-value searching in constant time. An Othello is collision-free and is significantly more compact than a traditional hash table as it does not store keys. But an Othello constructed on billions of *k*-mers still demands too much memory to be practical for use with standard computers. The hierarchical structure employed by SeqOthello overcomes this challenge using a divide-and-conquer approach. Specifically, *k*-mer occurrence maps are split into buckets according to their encoded lengths, with the assignment of each *k*-mer to its bucket determined by the root Othello. Within each bucket, the mapping between a *k*-mer and the location of its occurrence map is again stored in an Othello. SeqOthello significantly increases the volume of indexed *k*-mers within limited memory space and is inherently parallelizable.

**Figure 1.**
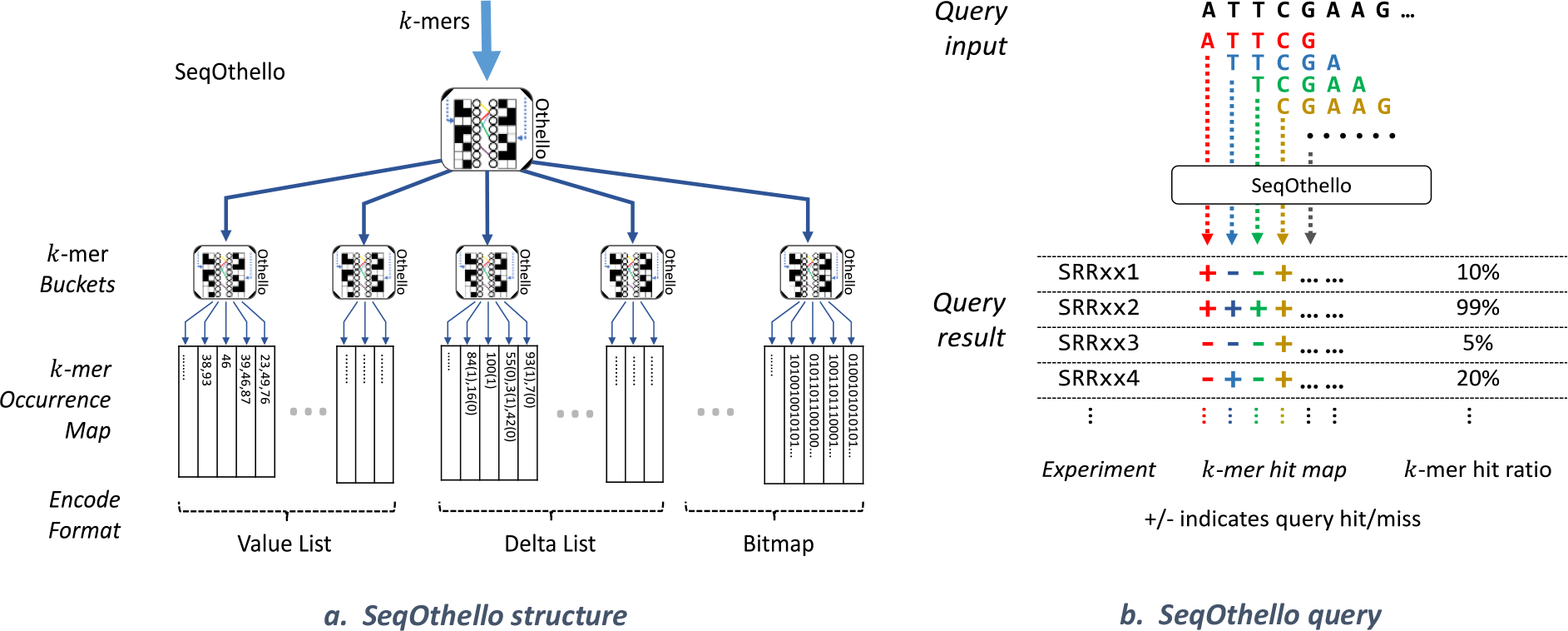
a. An illustration of the SeqOthello indexing structure to support scalable k-mer searching in large-scale sequencing experiments. The bottom level of SeqOthello stores the occurrence maps of individual k-mers, encoded in three different formats and divided into disjoint buckets. The mapping between a k-mer and its occurrence map is achieved by a hierarchy of Othello structures in which the root Othello maps a k-mer to its bucket and the Othello in each bucket maps a k-mer to its occurrence map. b. An example illustrating SeqOthello’s sequence-query process and output. A sequence query is decomposed into its constituent k-mers. The query result can be either a *k*-mer hit map, recording each k-mer’s presence/absence along the query sequence, *or k-mer hit ratios* (i.e., *the fraction of query k-mers present in each experiment*).

Querying of a SeqOthello first requires decomposing the query sequence into its constituent *k*-mers. The root Othello node identifies the occurrence bucket for each *k*-mer, following which each bucket Othello node retrieves the desired occurrence map. Per *k*-mer, this process requires exactly two Othello queries and is thus executed in constant time. The full set of occurrence maps is then synthesized to generate a *k*-mer hit map of the query for each experiment, where a hit means a k-mer is present in an experiment. Each *k*-mer hit map can be summarized into the number of hits, or a hit ratio, the fraction of hits out of the total *k*-mers in the query (Figure 1. b).

### SeqOthello outperforms State-of-The-Art algorithms

We compare SeqOthello to each of three state-of-the-art methods for querying large-scale RNA-seq datasets: SBT, SSBT, and SBT-AS. The evaluation was benchmarked on 2652 RNA-seq experiments of human blood, breast, and brain tissues from the SRA (Supp Table 1). We use Jellyfish^25^ to convert raw sequence data into *k*-mer files. Taking these files as input, SeqOthello requires 1.93 hours and a maximum of 14.1 GB memory to construct the index (Methods), 10 times faster than SBT and SSBT. At 20.8 GB, the SeqOthello index is 30% smaller than the most-compact SBT-based index, SSBT, and achieves a 700:1 compression ratio relative to the original database (Supp Table 2).

SeqOthello queries 198,093 transcripts from Gencode Release 25^26^ for *k*-mer hits in all 2652 experiments in 35.7 minutes using 15.2 GB memory. With four threads, the running time drops to 13.4 minutes. SBT-based queries only return the set of experiments whose *k*-mer hit ratio is greater than a user-defined threshold, denoted by *θ*. Even with a very high *k*-mer hit ratio (*θ* = 0.9), SBT-AS and SBT require 575 and 4,160 minutes to complete, respectively with higher memory cost than SeqOthello (Figure 2). While SSBT is extremely memory frugal, it is at the expense of much slower speed, two orders of magnitude slower than SeqOthello (Figure 2).

**Figure 2.**
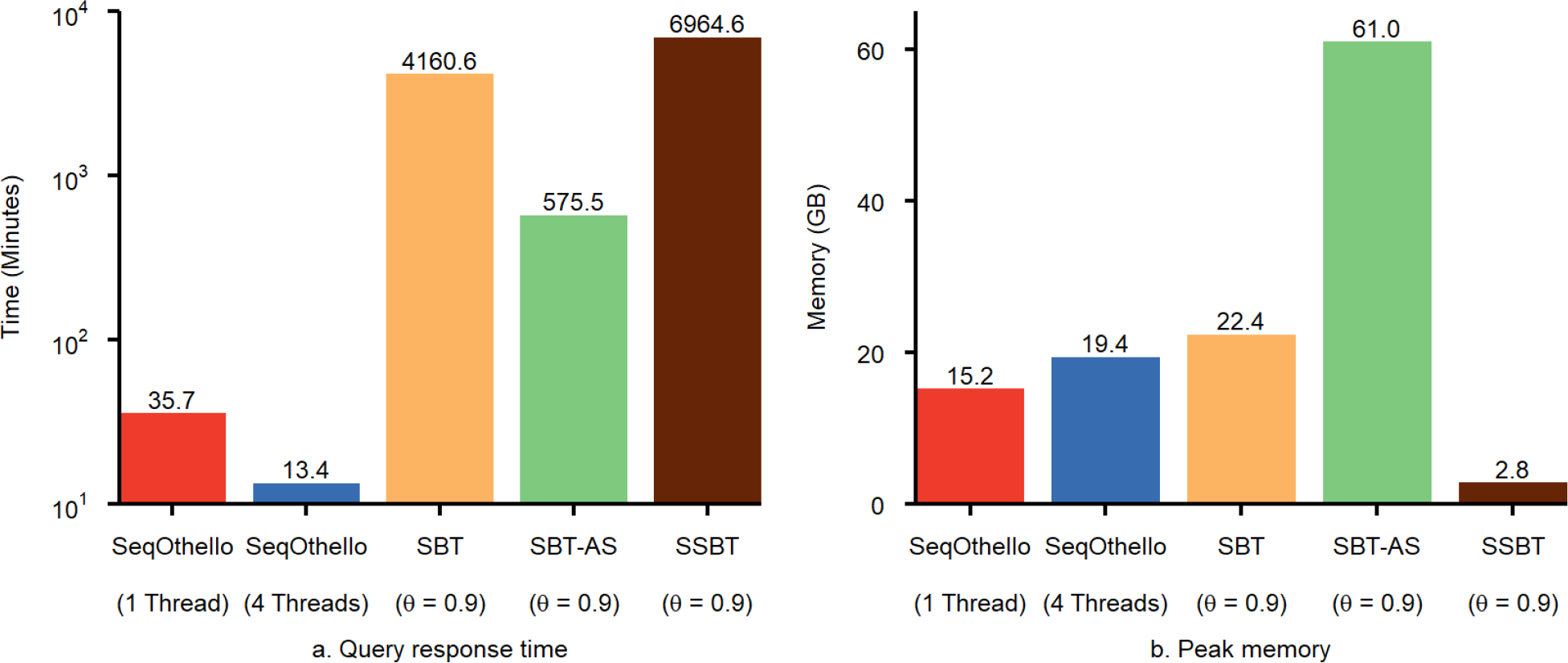
Performance comparison for querying with SeqOthello and each of three SBT-based algorithms: SBT, *SSBT*,and SBT-AS. Performance is benchmarked on 2652 human RNA-seq experiments. The query consists of 198,093 human transcripts in Gencode Release 25. a. Query response time. *b*. Peak memory.

The significance of experiments extracted by SBT using a single threshold *θ* is difficult to assess. To avoid generating misleading conclusions, multiple queries with different *θ* may be attempted to determine an approximate distribution, affording an overall query time several times larger than we report. Querying a small batch of 1000 transcripts with settings of *θ* = 0.7, *θ* = 0.8, and *θ* = 0.9 required 40 minutes to execute with SBT-AS, 190 minutes with SBT, and 241 minutes with SSBT (Supp Table 4). In contrast, SeqOthello requires only 4.6 minutes to query the same set of transcripts, and generates exact hit ratios for each transcript in each indexed experiment.

SeqOthello also accommodates online features for small-batch queries. Online queries preload the entire index into memory prior to querying, and can be executed in approximately 0.09 seconds per transcript (Supp Table 4). Our method’s advantageous speed permits it to support on-demand and instant queries from multiple users in a client-server setting. Other methods do not have online options at present.

### SeqOthello achieves near-exact k-mer query

To assess the accuracy in *k*-mer search, we queried 120,044,842 *k*-mers present in human transcriptome Gencode Release 25 against the SeqOthello index constructed for the aforementioned 2,652 experiments. We randomly selected 150 experiments and calculated the false-positive rate of *k*-mer queries in each experiment. The false positive rate is defined as the fraction of *k*-mers absent from the raw *k*-mer file that SeqOthello classifies as present with all queried *k*-mers. The Venn diagram (Supp Figure 2) shows an example of overlap among three sets of *k*-mers. SeqOthello recovers all *k*-mers that are truly present in the experiment, with guaranteed 100% recall rate. For *k*-mers that are not present in any of the indexed experiments, SeqOthello yields an extremely low rate of false positives: across 150 randomly chosen experiments, the average false-positive rate was only 0.015% with standard deviation of 0.071%.

To further evaluate the effect of false positives on transcript queries, we mapped the raw *k* -mers of each experiment to transcript sequences, calculating the true *k*-mer hit ratio for each transcript. We then compared the *k*-mer hit ratios generated by SeqOthello to the ground truth. Roughly 89.7% of transcripts afforded *k*-mer hit ratios equal to the true value, with an additional 9.3% exhibiting an error rate up to 0.003 (Figure 3). These results demonstrate that SeqOthello achieves near-exact query of *k*-mers and *k*-mer hit ratios. Additionally, as consecutive *k*–mers in a sequence are highly redundant, even a single base mismatch to the query sequence will be evidenced by the absence of multiple (i.e. *k*) *k* -mers, rendering an extremely low likelihood of false positive match due to alien *k*-mers (Methods). Although *k*-mer information is implicitly stored in bloom filters employed in SBT-based algorithms, efficient implementation of *k*-mer retrieval by these algorithms is not yet available.

**Figure 3.**
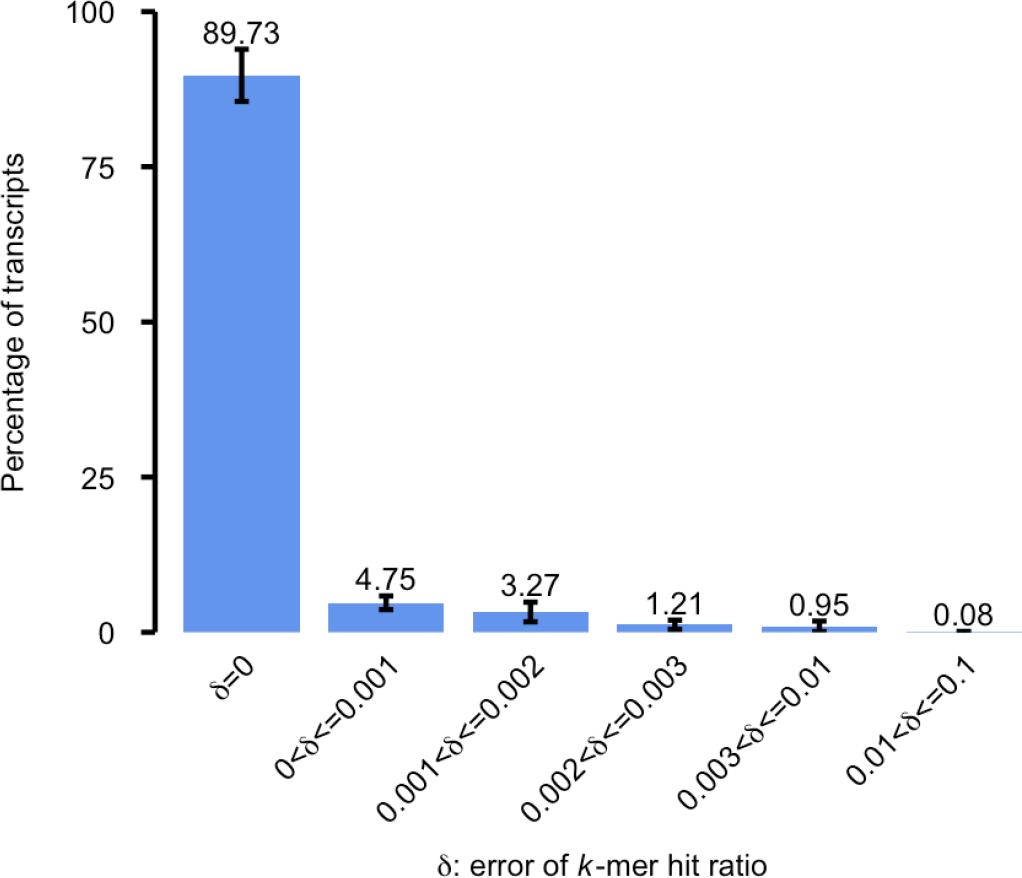
The distribution of error rate in k-mer hit ratios returned by SeqOthello. A randomly selected set of 150 experiments are extracted from SeqOthello’s result by querying all human transcripts on 2652 human experiments. The error (*S*) *of a transcript query over an experiment is calculated as the difference between the transcript’s k-mer hit ratio returned by SeqOthello and the k-mer hit ratio obtained by mapping raw k-mers using the same RNA-seq experiments to the transcript sequences. Each bar shows the percentage of transcripts with δ falls in a particular range. The error bar shows the standard deviation of such percentage measured on 150 experiments.*

### SeqOthello enables query against TCGA Pan-Cancer RNA-Seq Experiments

The Cancer Genome Atlas (TCGA)^2^ contains transcriptome profiles of 10,113 tumor samples obtained from 9,215 cancer patients. The database allows researchers to detect and characterize novel transcriptomic alterations across 29 different cancer types in the GDC Legacy Archive^2^. We have constructed a SeqOthello index, storing the occurrences of 1.47 billion 21-mers across all tumor samples. The preparation of *k*-mers averages 4 minutes per sample while the construction of SeqOthello on all samples took less than 9 hours. The index occupies only 76.6 GB of space, thus is portable for querying at different locations.

We apply the SeqOthello index to conduct a survey of all gene-fusion events curated by TCGA Fusion Gene Database as of December 2017^27^. The database documented of 11,658 unique tier-1 fusion events from TCGA detected by PRADA^28^. To query the presence of a fusion event, we construct a fusion junction sequence consisting of 20 bases from the donor exon and 20 bases from the acceptor exon, thereby guaranteeing that any 21-mer from the sequence will span the fusion junction (Supp Figure 3).

A SeqOthello query returns the number of *k*-mer hits on this fusion junction sequence in each sample. Determining a true fusion occurrence from *k*-mer hits is nontrivial. A simple approach specifies a minimum *k* -mer hit threshold. However, this technique yields lackluster sensitivity and specificity: Lowering the threshold permits detection with fewer spanning reads, but may increase false-positive calls in the presence of repetitive *k*-mers. Instead, we classify fusion events according to the distribution of *k*-mer hits in all tumor samples for each query. We first detect the background noise due to repetitive *k*-mers. Here the maximum number of repetitive *k*-mers is then estimated as the 98^th^ percentile in the distribution of *k*-mer hits, assuming less than 2% recurrence rate in all TCGA samples. (The highest recurrence rate yet documented is 0.953%, exhibited by TMPRSS2-ERG^27^.) We require a number of additional *k*-mers beyond this threshold as evidence of expression to conclude fusion in a sample (Supp Figure 4). Here 7 is chosen as it yields the optimal balance between sensitivity and specificity (Supp Figure 5).

Under this method, we detect 92.7% of tier-1 fusion occurrences in TCGA Fusion Gene Database^27^ with at least 10 spanning reads reported by PRADA. Additionally, we identify 270 novel occurrences of fusion events across 17 tumor subtypes that are not identified by PRADA. We selected two fusion pairs with occurrences most inconsistent with current curation for further validations: FGFR3-TACC3 in GBM samples (5 novel, 3 undetected) and ESR1-C6orf97 in BRCA samples (2 novel, 5 undetected). We were able to confirm all 7 novel fusion occurrences by the identification of at least 10 fusion spanning reads supporting each. For all undetected fusions, insufficient spanning reads were confirmed, which are consistent with low read support recorded in the database (Supp Files).

Figure 4 depicts the 10 novel, recurring fusions with greatest number of occurrences suggested by SeqOthello. Quite a few have doubled or even tripled the original recurring rates. Interestingly, all novel occurrences agree with the original fusion cancer-type classifications, rendering the chance of random occurrence negligible. This result corroborates their cancer specificity and supports the high precision of SeqOthello’s query results. One example of this consistency is TMPRSS2-ERG, a clinical marker for prostate cancer. SeqOthello extracted 122 pre-identified occurrences of TMPRSS2-ERG and 142 novel occurrences, all from prostate cancer samples. The complete information of all detected fusion occurrences is listed in Supp Files.

**Figure 4.**
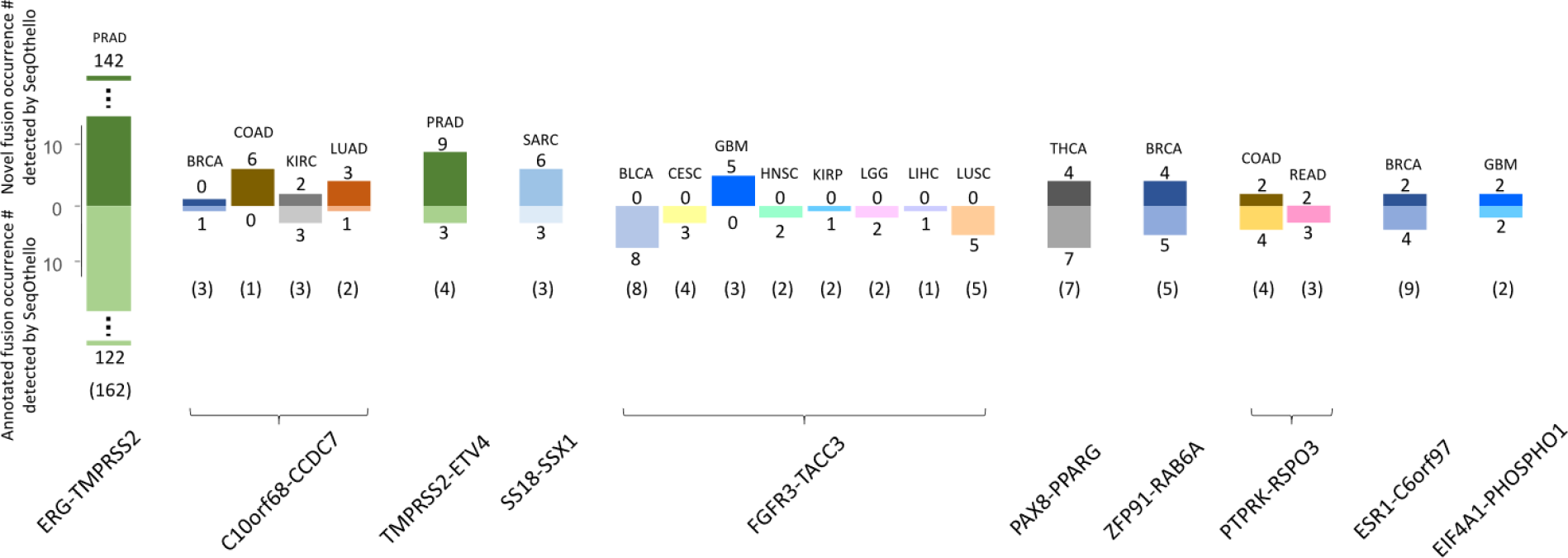
Ten fusion gene pairs with the highest novel occurrences identified by SeqOthello. We used SeqOthello to conduct a survey of all fusions curated in TCGA gene-fusion database against 10,*113 TCGA Pan Cancer RNA-seq samples. Each column in the bar plot represents fusion occurrence in a cancer type. The number in parenthesis below each bar indicates the number of documented occurrences reported in the current database.*

## Discussion

SeqOthello is a novel algorithm capable of indexing large-scale RNA-seq experiments that supports online sequence query. We constructed a SeqOthello index on the TCGA Pan-Cancer RNA-seq datasets, the latter totaling 54 TB in compressed fastq format. The SeqOthello index uses only 76.6 GB disk space, achieving a compression ratio of 700:1. Querying the index to assess the prevalence of 11,658 documented fusion events requires only five minutes on a standard desktop computer with 32 GB memory. This performance is orders of magnitude faster than the most-efficient existing fusion-detection algorithm, estimated to require 785 days of computation to process all TCGA data (methods).

SeqOthello queries on individual *k* -mers for their presence and absence in each experiment. This approach enables access to sequencing-level information without downloading and reanalyzing raw reads. For example, *k*-mers unique to a transcript isoform can be used to distinguish alternative isoforms (Supp Figure 7). Additionally, SeqOthello enables large-scale integrative analysis by querying. This functionality is evidenced by our fusion-calling method, which leverages the global distribution of *k*-mer hits queried with SeqOthello to achieve accurate fusion classification.

The simple query supported by SeqOthello is powerful, with myriad applications yet to be defined. One can use SeqOthello to assess the prevalence of clinically important features in different patient populations or to compare across different patient cohorts. Beyond transcripts, one can use SeqOthello to identify expressed regions by querying entire reference genomes. SeqOthello can be potentially leveraged on any form of next-generation sequencing data that can be translated to a *k*-mer occurrence matrix. We leave the definitions and demonstrations of these applications for future work.

In conclusion, SeqOthello is parameter-free, reference-free, and annotation-free. Its unbiased nature supports large-scale integrative and comparative studies, while its ultra-fast performance and undemanding system requirement render it appropriate for a wide variety of research investigators. SeqOthello will enable novel discoveries that would be otherwise unrealizable for individual research labs.

## Methods

### Section 1. The Othello data structure

The mapping of *k*-mers in either level of SeqOthello is maintained by a data structure named *Othello*. Othello belongs to the class of minimal perfect hashing algorithms^29^ and has demonstrated superb scalability of both memory and querying speed in various applications^23,24,30,31^. Othello implements a hashing classifier to efficiently map keys (*k*-mers) to appropriate categories.

An Othello ***O***(*S,V*) maps a predefined set of *k*-mers *S* to a list of categories represented as integers, denoted by *V* = {1,2,…,*v*}. Let *T:S* →*V* be the function that maps *k*-mers in *S* to classes in *V*, where *T*(*s*) indicates the category of a *k*-mer *s* ∈*S.* Each category in *V* is represented by an *l*-bit integer, where *l* = [log_2_(*v*+ 1)].

In essence, an Othello ***O***(*S,V*) maintains a query function τ*:U* → *C* mapping the set of all possible *k*-mers, *U*, to the set of all *l*-bit integers, *C* = {0,1,…,2^l^− 1}. Thus *S* ⊂*U* and *V* ⊂. Furthermore, τ is a superset of *T*: That is, for any *s* ∈*S*,τ(*s*)*= T*(*s*); for any *s’*∈ *U* − *S*,τ (*s*) is a deterministic *l*-bit integer. A *k*-mer *s’* is called ***alien*** if and only if *s’*∈*U* – *S*, such that *s’*∉*S* and the mapping for ***s’*** is not specified in ***T.***

#### Section 1.1 Properties of Othello

We described previously the Othello data structure in MetaOthello ^23^ (Section 2.2.1, Fig.2). We summarize the properties of Othello as follows.

- An Othello data structure maintains the mapping τ*:U* → *C*, where *C* = {0,1,…,2^*l*^– 1} and *l* =[log_2_*v*+ 1].
- Implementation of Othello entails (1) a pair of hash functions, ⟨*h_a_,h_b_*⟩, and (2) two arrays of *l*-bit integers, *A* and *B*. The lengths of the arrays, respectively denoted *m_a_* and *m*_b_, satisfy 2.67*n* ≤ m_a_+ m_b_< 4*n*, where *n* is the number of *k*-mers. The functions and contents of the arrays are determined by the construction algorithm according to the keys and their corresponding categories. The time complexity of the construction algorithm is *O*(*n*).
- An Othello built to map *nk*-mers to *v* categories requires at most 4*n*[log_2_(*v*+1)] bits of memory space.
- Given a *k*-mer s, its class information τ(*s*) is computed by [*h_A_* (s)] ⊕*B*[*h_B_*(*s*)]. Thus, querying a *k*-mer requires only two memory accesses and one XOR bit operation, making it extremely fast.

#### Section 1.2 On alien k-mer query of Othello

Let τ(s) be the category returned by querying a *k*-mer s on Othello ***O***(*S,V*). If *s* ∈*S*, then Othello guarantees that τ(*s*) = *T*(*s*). An alien *k*-mer *s*′∉*S* may be correctly recognized as an alien if τ(*s*′)∈ *C* − *V*; alternately, such a query may return a false positive if τ(*s*′)∈ *V.* Our next goal is to analyze and bound the probability of Othello in recognizing alien *k*-mers.

##### Lemma 1

For any alien *k*-mer *s*′ ∉*S*, a query on Othello ***O***(*S,V*) returns an *l*-bit integer τ(*s*′). For any integer *x*, the probability of τ(*s*′) = *x* is denoted by *p_x_* 
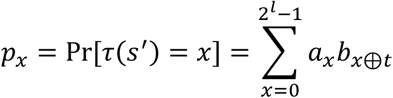

**Proof:** Here *a_x_* is the fraction of 0s in the array *A* of the Othello data structure, and *b_x_* is the fraction of 0s in array *B*. The values of *a_x_* and *b_x_* are computed using the content stored in the memory of Othello. Lemma 1 is a direct application of a result presented in our previous work (MetaOthello ^3^, Section 2.2.3).

##### Lemma 2

Let |*S*| = *n* for an Othello ***O***(*S,V*) constructed with *n* elements. Then *p_0_>* 0.223 as *n*→∞.

**Proof:** We prove Lemma 2 by giving an estimated lower bound on *p_0_*, which is the probability of an alien *k*-mer being assigned to category 0 (one of the alien categories). Array *A* of the Othello contains *m_a_* elements. Each *k*-mer is mapped to an index of array *A* computed by *h_a_*(*s*), where *h_a_* is a uniform random hash function. Assuming the number of *k*-mers, *n*, is large, the possibility of an index in *A* not being hit by any of the *h_a_*(*s*) values is

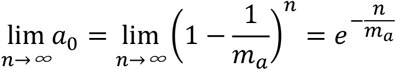

An analogous statement holds for array *B*. Note that 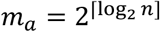 and 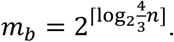 We have

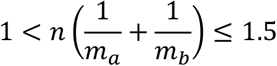

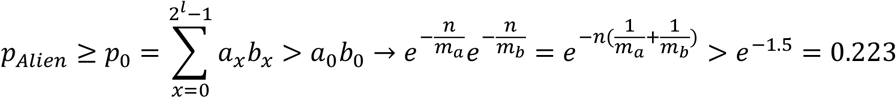

##### Theorem 1

For any alien *k*-mer *s′*∉*S*, the probability that *s’* is identified as ‘alien’ by an Othello ***O***(*S,V*) is given by:

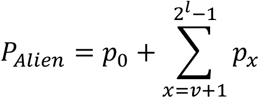

We also have *p_Alien_* > 0.223 as |*S*| → ∞.

**Proof:** The probability that an alien *k*-mer falls into a class *x* ∈ *C* − *V*, denoted *p**_x_***, can be computed using the approach specified in Lemma 1. Note that *C*−*V* = {0,*v*+ 1,*v* + 2, …,2*^l^*− 1}, so that:

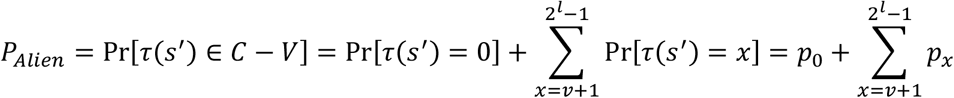

By Lemma 2, *p_Alien_* ≥ *p_0_* > 0.223 as |S| → ∞.

## Section 2. Encoding of k-mer occurrence map

We define the occurrence map of a *k*-mer as a binary vector recording the *k*-mer’s presence or absence in each experiment. Given *m* experiments, the occurrence map can be stored using *m* bits, where 1 represents presence and 0 represents absence in a certain experiment. To minimize the storage requirement of these vectors, we have developed a hybrid encoding method that leverages one of three different encoding strategies depending on the occurrence frequency of a *k*-mer. Each *k*-mer is stored using the method that yields the shortest code. These encoding methods are detailed below:

- Value-list encoding. This method is used to compress occurrence maps associated with rare *k*-mers. For an *m*-bit occurrence map with exactly *t* 1s (representing presence in *t* out of *m* samples), we enumerate the *t* indices of these positions as a list. Each index is represented by *t* integers, each [log_2_*m*] bits long. This list can also be viewed as a *t*[log_2_*m*]-bit integer. Value-list encoding is used when *t*[log_2_m]≤ 64.
- Delta-list encoding. This approach is employed for occurrence maps with a relatively larger number of 1s (*t*[log_2_*m*] > 64). The *m* elements in the occurrence map can be considered as a succession of alternating subsequences of 0s and 1s. Thus the map can be represented by a list of 2*w*+ 1 integers, ⟨*x_1_,y_1_,x*_2_,*y_2_*,…,*x_w_,y_w_,x_w+1_*⟩, representing the number of digits in each subsequence, where *x_1_* ≥ 0, *x_w+1_* ≥ 0; *y_1_,y_2_*,…,*y_w_* ≥ 1,*x_2_,x_3_*,· ··,*x_w_* ≥ 1; and *x_1_* + *y_1_+ x_2_* + *y_2_*+ … + *x_w_*+ *y_w_*+ *x_w +1_*=*m*. The occurrence map can be reconstituted by enumerating *x_1_* 0s, followed by *y_1_* 1s, *x_2_* 0s, *y_2_* 1s, etc. For example, consider an occurrence map of *m* = 20 elements, 1110011…10, with 1s at indices 1,2,3,6,8,9,…,19. The corresponding delta-list representation is ⟨*x_1_* = 0,*y_1_* = 3,*x_2_* = 2,*y_2_* = 14, *x_3_* =1⟩. The 2*w* + 1 integers from this first step are further encoded as positive integers. Multiple procedures exist for the second encoding step, the choice of which depends on the relative importance of minimizing encoding/decoding overhead *versus* maximizing the compression rate. To balance the time and memory complexity of encoding, as well as the storage overhead, we choose to encode the delta list as a hexadecimal stream. Each integer is converted to a hexadecimal value using the method described in Method Table 1. We then concatenate the hexadecimal values into a single hexadecimal datum. For the delta list shown in the example, ⟨ 0,3,2,14,1⟩, the corresponding hexadecimal format is 0×8, 0×B, 0×A, 0×4E, 0×9. After concatenation, the final result is 0×8BA4E9.
- Bitmap encoding. Each occurrence bitmap is an *m*-bit value, with each bit coding the presence or absence information for one of the *m* samples. As this method requires more memory than other options, it is used only when other options do not work.

**Method Table 1.**
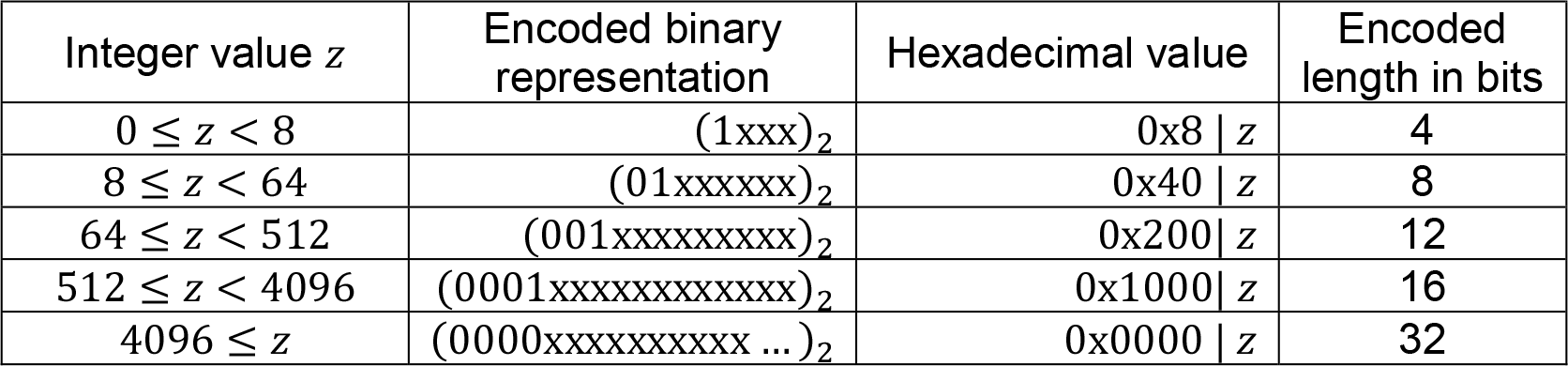
Hexadecimal encoding for integer values in delta-list encoding

## Section 3. Construction of SeqOthello

### Section 3.1 Construction algorithm

Construction of a SeqOthello data structure requires as input a list of *k*-mer files, each containing the set of *k*-mers extracted from reads associated with a distinct RNA-seq experiment. Currently the *k*-mer file is generated by applying Jellyfish to fastq files.

**Step 1: Assembling the occurrence map of each *k*-mer in the collection of experiments to be indexed.** The goal of step 1 is to determine each *k*-mer’s presence/absence information across all experiments. This task requires the integration of *k*-mers from all *k*-mer files, but simultaneous file access is time-consuming and not allowed by many operating systems. Instead, we employ a strategy similar to merge sort. We first obtain *k*-mer occurrence maps for small groups of experiments, where each group contains approximately 50 samples. These intermediate occurrence maps are encoded as delta lists, which significantly reduces file sizes. The groups are then merged to obtain the ***k***-mer occurrences across all experiments. After SeqOthello is constructed, the group files generated at this step are no longer needed. However, as these files are orders of magnitude smaller than the original *k*-mer files, they can be stored to support update of the SeqOthello structure.

**Step 2: Assignment of *k*-mer occurrence maps to buckets.** We next divide the entire set of *k*-mers into disjoint buckets based on their occurrence maps using the following principles: (1) Occurrence maps within the same bucket should be generated by the same encoding approach; (2) the lengths of encoded occurrence maps within the same bucket should have limited variation; and (3) the total size of the encoded occurrence maps within each bucket should not exceed a specified threshold (by default, 128 MB).

Given a maximum bucket size, we define the range of encoding lengths for each bucket prior to allocating *k*-mers. Note that the distribution of *k*-mer encoding lengths is unknown prior to construction. To avoid multiple iterations over all *k*-mers during bucket assignment, we designed a sampling-based approach to estimate the range of encoding lengths. The goal is to set an open upper bound *n*_*t*+1_ and closed lower bound *n_t_* so that *k*-mers with encoding lengths in the range [*n_t_,n*_*t*+1_) are assigned to each bucket *t*. We select 10 million *k*-mers, which is approximately 0.1% of the *k*-mers present over all experiments, and let *L_i_* be the estimated number of *k*-mers with encoding length equal to *i*. Starting from *t* = 1 and *n_1_* = 1, we greedily select the maximum index *n*_*t*+1_ so that 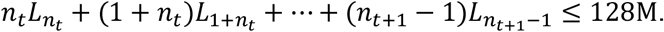 Once the number of buckets and their ranges of encoding lengths are determined, the construction algorithm will iterate over each *k*-mer, assigning it to the appropriate bucket in accordance with the encoding length of its occurrence map. The encoded occurrence maps are further compressed by gzip when the final structure is stored as a file.

**Step 3: Establish *k*-mer mapping using Othello.** During step 2, SeqOthello maintains the list of *k*-mers and their corresponding encoded occurrence maps in each bucket. Once the *k*-mer assignment is completed in the bucket, an Othello will be established to record the mapping between *k*-mers and the locations of their occurrence maps. Once the buckets are finalized, a root Othello is constructed to record the mapping between the entire set of *k*-mers and their bucket IDs.

SeqOthello also maintains an .xml file to store metadata associated with the data structure, which includes basic information about the experiments and information necessary for the query algorithm to interpret the data file.

### Section 3.2 Optimization for k-mers that appear in only one experiment

The prevalence of individual *k*-mers varies dramatically, with plots often exhibiting a *U*- or *L*- shaped distribution (Supp Fig. 1). Note that the number of *k*-mers present in only one experiment is relatively large compared to *k*-mers with higher frequencies. We apply the following approach to improve the efficiency and accuracy of SeqOthello.

Instead of storing all *k*-mers with single occurrence in a level-2 bucket, we encode them directly in the root Othello. Let *E* be the set of experiments indexed by SeqOthello, identified by integers {1,2, …,|*E*|}. Let *B* be the set of buckets identified by integers {|*E*| + 1, |*E*| + 2, …,|*E*| + |*B*|}. The root Othello records the mapping between *k*-mer set *S* and *E*∪ *B*. For any *k*-mer *s*, if the query result on the first level τ(*s*) ∈ {1,2, …|*E*|}, SeqOthello will report that *s* is present in the experiment with index τ(*s*); if τ(*s*) = |*E*| + *b* for some integer *b* ∈ {1,2,…, | *B*|}, then τ(*s*) ∈*B* and the query process will continue into the bucket with index *b* on the bottom layer of SeqOthello.

### Section 3.3 Insertion of new experiments into SeqOthello

If the group files generated at Step 1 have been retained, the insertion of new experiments to SeqOthello is quite fast, especially for batch update. The process involves merging newly inserted experiments with the existing group files, and then repeating Steps 2 and 3 of the above construction algorithm. The entire update requires only a few hours to complete.

## Section 4. The probability of false-positive k-mer query with SeqOthello

SeqOthello maintains a mapping from a large set of *k*-mers to their occurrence maps. However, due to the nature of Othello being a minimal perfect hashing classifier, querying of an alien *k*-mer (*i.e.,k*-mer that does not exist in any of the samples) with SeqOthello may afford a false report of its presence in one or more RNA-seq experiments. Here, we analyze the likelihood of such a false report.

### Section 4.1 Notations

In reference to SeqOthello, we use the notation *^Root^****O***(*S,V*) to denote the root-level Othello. *^Root^****O***(*S,V*) records the mapping between a *k*-mer in *S* and its assignment either to a single experiment or to a second-level bucket in *V* = *E* ∪ *B*.

For any bucket *b* ∈ *B*, we use the notation *^b^* ***O***(*S_b_,V_b_*) to denote the associated Othello, where *^b^* ***O***(*S_b_,V_b_*) stores the mapping between a *k*-mer in *S_b_* and its occurrence map index in *V_b_.* Thus *S_b_* is the set of *k*-mers that are assigned to bucket *b* and *V_b_* = {1,2,…,*v_b_*} is the list of indices for encoded occurrence maps in bucket *b*.

We list the primary notation used in the following analysis in Method Table 2.

**Method Table 2.** A summary of notations used in Section 4

|  |  |
| --- | --- |
| $^{Root}\mathbf{O}(S, V)$ | Othello at the root of SeqOthello |
| $^b\mathbf{O}(S_b, V_b)$ | Othello of the bucket $b$ |
| $E$ | Set of RNA-seq experiments |
| $B$ | Set of buckets |
| $W_t$ | $t^{th}$ occurrence map in a bucket $b$ |
| $^{SeqOthello}P_{Alien}$ | Probability of an alien $k$-mer being recognized as Alien by SeqOthello |
| $^{SeqOthello}P(e)$ | Probability of an alien query returning experiment $e$ |
| $^{root}p_x$ | Probability that query of an alien $k$-mer on the root Othello $\tau(s')$ returns $x$ |
| $^bp_x$ | Probability that query of an alien $k$-mer on the Othello in bucket $b$ returns $^b\tau(s')$ value $x$ |

### Section 4.2 Probability of alien k-mer recognition and false positive presence

Let *s′* be an alien *k*-mer, and τ(*s′*) be the result returned when querying *s′* on the root Othello. Then τ(*s’*) falls into one of the following three categories:

A. τ(*s’*)∉*V*, where *V* = *E* ∪ *B*. This *k*-mer will be identified as alien, and SeqOthello will report its absence from the database. The probability of this result is *^root^P_Alien_*, which can be calculated according to Theorem 1.
B. τ (*s′*) ∈ *E*. Such a *k*-mer will be reported falsely as existing in the experiment identified by τ(*s′*). For any experiment *e* ∈*E*, the probability of returning *e* as the result of querying an alien *k*-mer has a probability *^root^p_e_*, which can be calculated based on Lemma 1.
C. τ(*s′*) ∈ *B*. In this case, the query process would continue into the bucket *b* identified by τ(*s′*).This circumstance occurs with probability *^root^ p*_|*E*| + *b*_. Inside the bucket *b*, the query *^b^*τ(*s′*) will result in one of two scenarios:

1. *^b^*τ(*s′*) ∉ *V_b_*. In this case, *s′* is identified as alien in bucket *b* with probability *^b^P_Alien_*, which is *P_Alien_*for the Othello *^b^**O***(*S*_*b*_,*V*_*b*_).
2. *^b^*τ(*s′*) ∈ *V_b_*. Here *s′* is mapped falsely to a location storing the occurrence map of a different *k*-mer. A calculation follows for the probability of this outcome. Assume there are *v_b_* encoded occurrence maps stored in bucket *b*, namely *W_1_,W_2_*, …,*W_vb_*. We use the notation *W_t,_* ∈ {0,1} to denote the presence/absence information for experiment *e* stored in the *t*-th occurrence map. Here, *W*_*t,e*_ = 1 indicates that the *k*-mer associated with occurrence map *W_t_* is marked as ‘present’ in experiment **e* W_t_*,_*e*_ = 0 indicates it is marked as ‘not present’ in experiment *e*. Note that a query on bucket *b* returns the occurrence map with index *^b^*τ(*s′*), namely **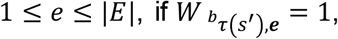** For any experiment *e*,1 ≤ *e* ≤ |*E*| if **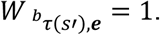** then the query result would indicate falsely that *s′* presents in experiment *e*. We use the notation *^b^P*(*e*) to denote the probability of the query on bucket *b* yielding **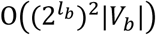**. *^b^P*(*e*) is equal the probability of *^b^*τ(*s′*) returning any index *x* such that the *x*-th occurrence map *W_x_* satisfies *W_x_*,_*e*_ =1:

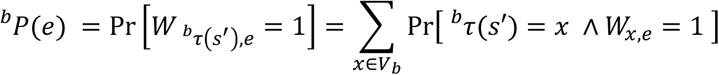 Noting that *W_x,_*∈ {0,1}, 
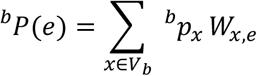 Computing *^b^p_x_* for all *x* ∈ *V_b_* using Lemma 1 requires 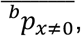 computation, which becomes infeasible when *l_b_* is large. Hence, we use an alternative approach to estimate the *^b^p_x_* values when *l*≥ 12. Lemma 2 indicates that the value of *^b^p_0_* is significantly larger than *^b^p_x_* values for *x* ≠ 0. We also observe that the values for *^b^p_x_* are similar for any *x* ≠ 0 and *x*< 2*^l^_b_* in the same bucket *b.* We therefore use the average value of *^b^p_x_* over *x* ≠ 0, denoted by **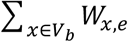** to replace individual *^b^p_x_* values: 
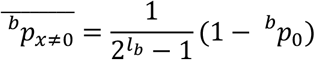 Hence, 
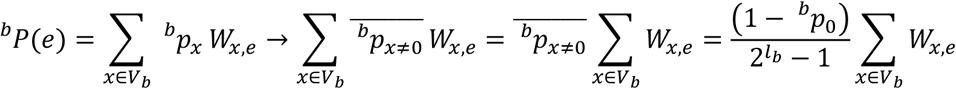 Here, **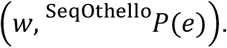** the number of encoded occurrence maps in bucket *b* in which the associated *k*-mer is marked to be present in experiment *e*. For an alien *k*-mer *s*′, the query on SeqOthello may return a false presence in experiment *e* if τ(*s′*)falls in category B, a circumstance which occurs with probability *^root^p_e_*. Otherwise, if τ(*s’*)satisfies circumstance C.2, the query yields an occurrence map in which experiment *e* is marked as positive with probability *^b^P*(*e*). Hence, the probability of an alien *k*-mer query on the two-level SeqOthello yielding a false-positive presence in experiment *e* is:

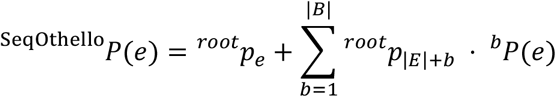 On the other hand, an alien *k*-mer has a very good likelihood of being recognized as alien if τ(*s′*) satisfies circumstance A, or falls in circumstance C and is subsequently identified under C.1. Taken together, the overall probability of SeqOthello identifying the *k*-mer as alien is:

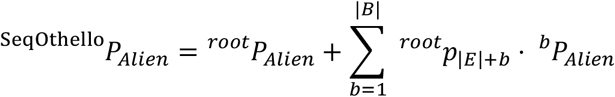 We present a numerical estimation of various probabilities based on the distribution of k-mer occurrences as well as the SeqOthello structures constructed for the two datasets used in this paper. The results are given in Method Table 3 below.

**Method Table 3.**
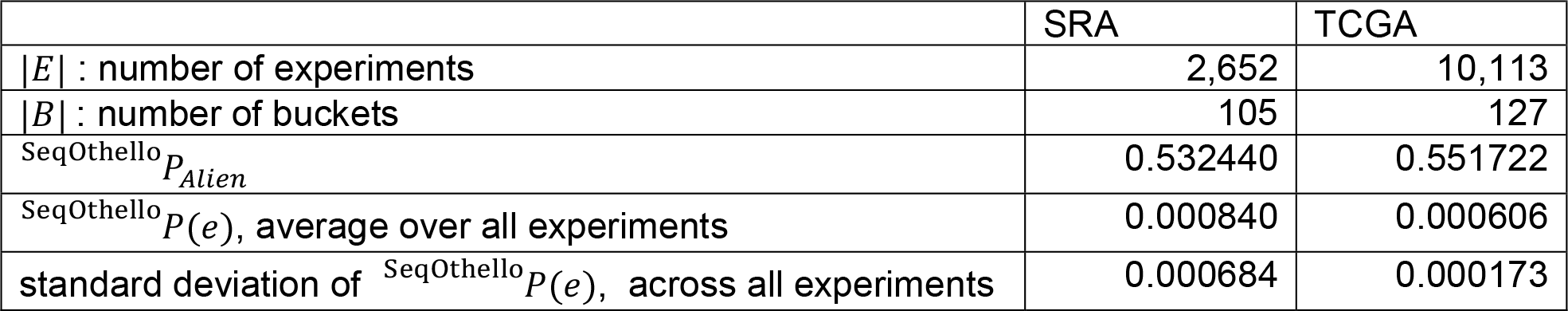
Estimated probability values computed on SeqOthello constructed for Human and TCGA datasets

### Section 4.4 Error rate of a SeqOthello sequence query

SeqOthello executes sequence query by making individual *k*-mer queries extracted from the sequence. The probability of returning false-positive *k*-mer hits is low and can be computed as *^SeqOthello^P*(*e*). Let *X*(*e*) be the number of false positives for experiment *e* returned over *w* alien *k*-mer queries. Then, *X*(*e*)follows the binomial distribution **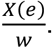** Note that the query result for transcript query is reported as the fraction of present *k*-mers for each sample, and *X*(*e*) false positive *k*-mers will result in an error rate of 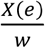. Note that the 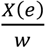 is usually 0. The probability of 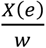 being large enough to affect the query result is very low, only occurring when multiple *k*-mer queries return the same false-positive experiments. For example, for *w* = 50 and *P*(*e*) = 0.0084, the probability of *X*(*e*) > 2 is 1.15×10^−5^. Thus, SeqOthello returns the query result with error rate 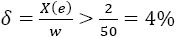 with probability 1.15×10^−5^,which is much lower than the probability of a single error.

## Section 5. Performance comparison

### Section 5.1 System Configuration

All comparison tests with SBT, SSBT and SBT-AS were conducted on a Linux OS (RHEL) sever with Quad Intel E5-4640 8 core (Sandy Bridge) @ 2.4 GHz processors, 512 GB of 1600 Mhz RAM, and 4 x 1 TB local (internal) NLSAS disk.

### Section 5.2 Versions and parameters for SBT, SSBT, SBT-AS

SBT, SSBT, and SBT-AS versions used in the evaluation are provided in Method Table 4.

**Method Table 4.** SBT, SSBT, and SBT-AS version information

| Algorithm | URL | Version |
| --- | --- | --- |
| SBT | https://github.com/Kingsford-Group/bloomtree | f7986e4511189cb781b4e3517626b396fb11eefa |
| SSBT | https://github.com/Kingsford-Group/splitsbt | 0fe43f4c0de7a0a452486a252a5e317862c2af45 |
| SBT-AS | https://github.com/medvedevgroup/bloomtree-allsome | 383d23f17d5537a0abf1436e5d04795ef91950b3 |

## Section 6. Estimation of typical fusion-detection processing time

A study published recently by Kumal et al (2016) details a comprehensive comparison of 12 fusion-detection algorithms, including FusionHunter ^32^, FusionMap^33^, Bellerophontes^34^, MapSplice ^35^,Chimerascan^36^, TopHat-Fusion ^37^, BreakFusion ^38^, SOAPfuse ^39^, JAFFA ^40^, nFuse ^41^, EricScript ^42,^and FusionCatcher ^43^. The authors reported that it requires 120 to 3,845 minutes for current tools to process a dataset of 70 million paired-end reads, averaging between 1.71 and 54 minutes per million reads. The TCGA RNA-seq dataset is estimated to contain 660 billion reads. Taking the fastest processing speed regardless of accuracy, we estimate that it costs 785 days of computation to process all the TCGA data for fusion detection using standard tools.

### Section 7. Code availability

The source code of SeqOthello and the scripts used to build and query the SeqOthello mapping are available at https://github.com/LiuBioinfo/SeqOthello.

---

**Supplementary Figure 1.**
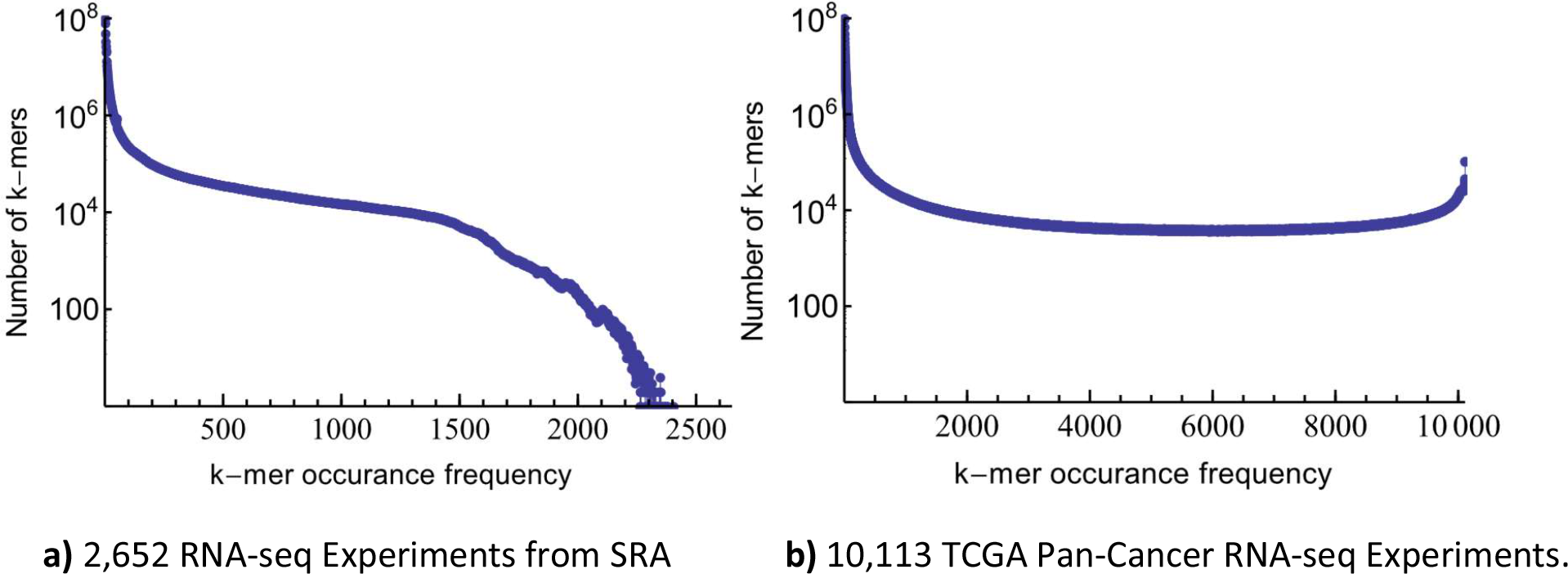
The histograms of *k*-mer occurrence frequencies in two human RNA-Seq datasets. A *k*-mer’s occurrence frequency is the number of samples containing the *k*-mer. The difference in the high frequency *k*-mers between the two datasets suggests less homogeneity in RNA-seq experiments downloaded from SRA than these generated by TCGA. **a)** The *k*-mer occurrence histogram across 2652 RNA-seq experiments of human blood, breast and brain tissues from the SRA. **b)** The *k*-mer occurrence histogram across 10,113 TCGA Pan-Cancer RNA-seq experiments.

**Supplementary Figure 2.**
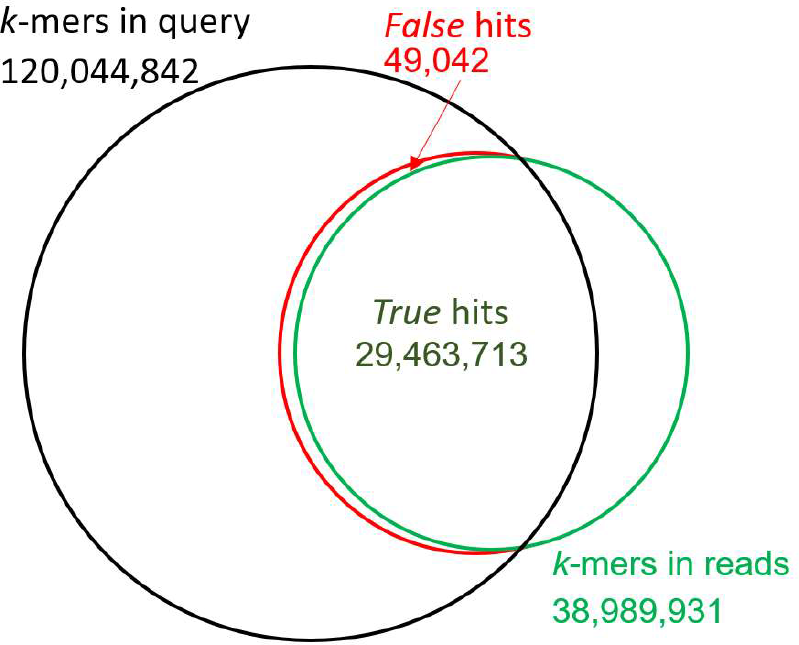
A Venn-Diagram showing the accuracy in querying human transcriptomic ***k-*** mers totaling 120,044,842 from experiment SRR925711. There are 120 million transcriptomic *k*-mers used for the SeqOthello query (the dark circle) and within the particular experiment SRR925711, there exists 39 million *k*-mers (the green circle). SeqOthello returned a total of 29.5 million hits in the experiment, among which only less than 0.2% is false. This is comparable to error rates found in next-generation RNA-seq data. Similar exercises were repeated on 150 experiments to obtain a representative distribution of false positive rate in *k*-mer query. (Figure 3)

**Supplementary Figure 3.**
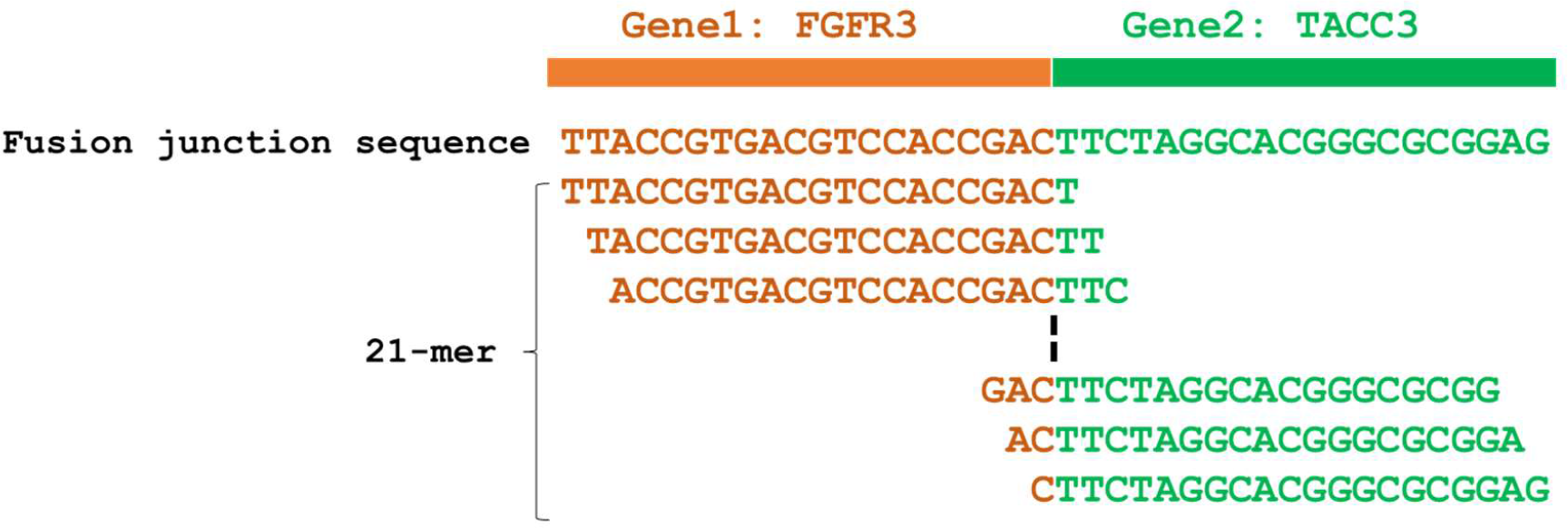
An illustration of fusion junction sequence constructed for fusion query using SeqOthello. Each fusion junction sequence consists of 20 bases from donor exon in one gene and 20 bases from acceptor exon in the other gene. Each 21-mer within this 40-base sequence spans the fusion junction. The query of a fusion sequence using SeqOthello may return a maximum of 20 *k*-mer hits for each RNA-seq Experiments indexed by SeqOthello.

**Supplementary Figure 4.**
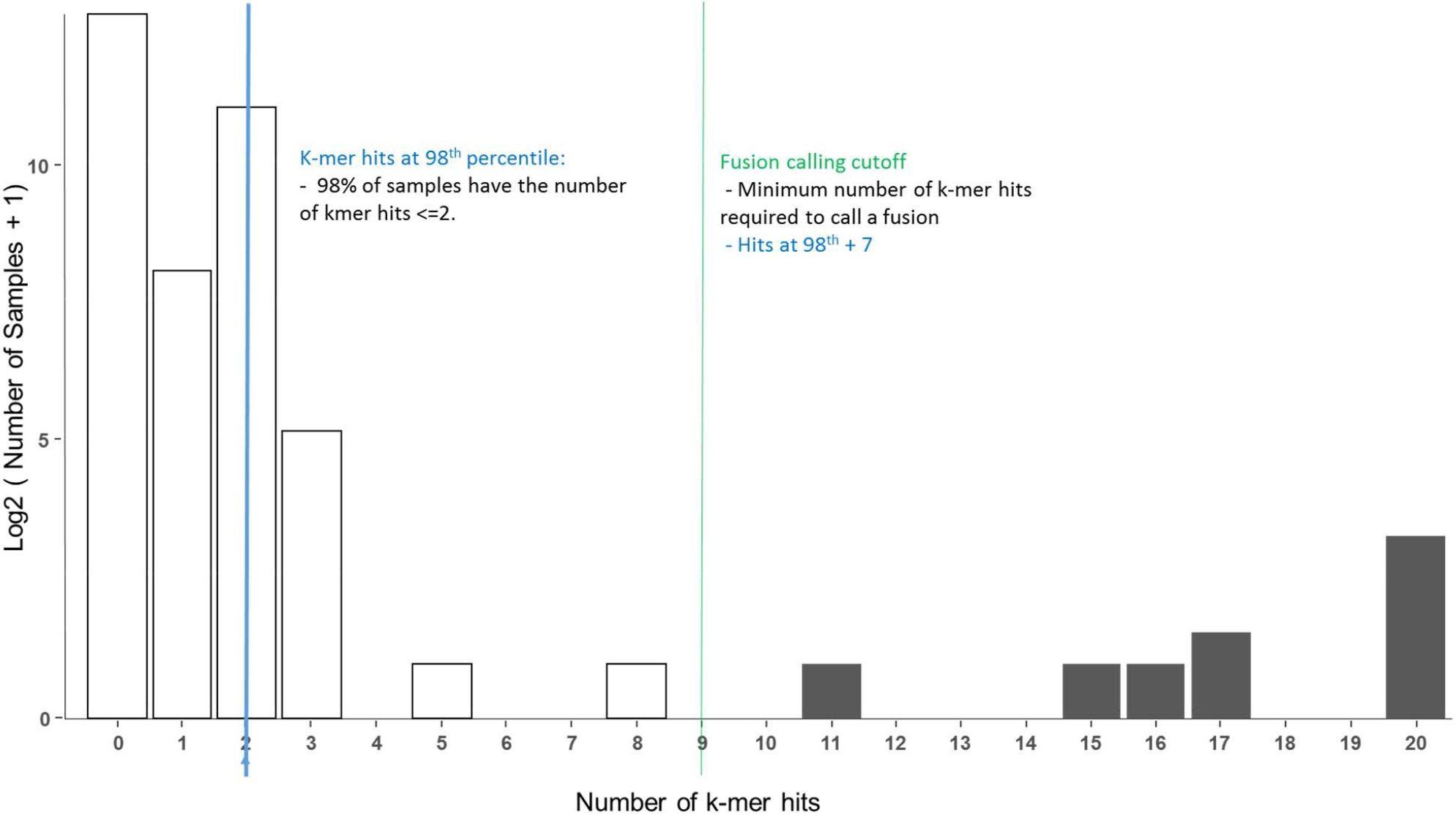
An illustration of SeqOthello gene fusion calling method based on k-mer hit distribution. Shown in the figure is the distribution of the number of *k*-mer hits when querying fusion event TMPRSS2-ERG (chr21: 42880008 – chr21: 39956869). We identify the number of *k*-mer hits at 98^th^ percentile, which is 2 in this case. The fusion calling cutoff requires 7 additional *k*-mers for the evidence of expression (According to information in Supplementary Figure 5). For this particular fusion event, a fusion is called if the experiment has more than 2+7=9 *k*-mer hits.

**Supplementary Figure 5.**
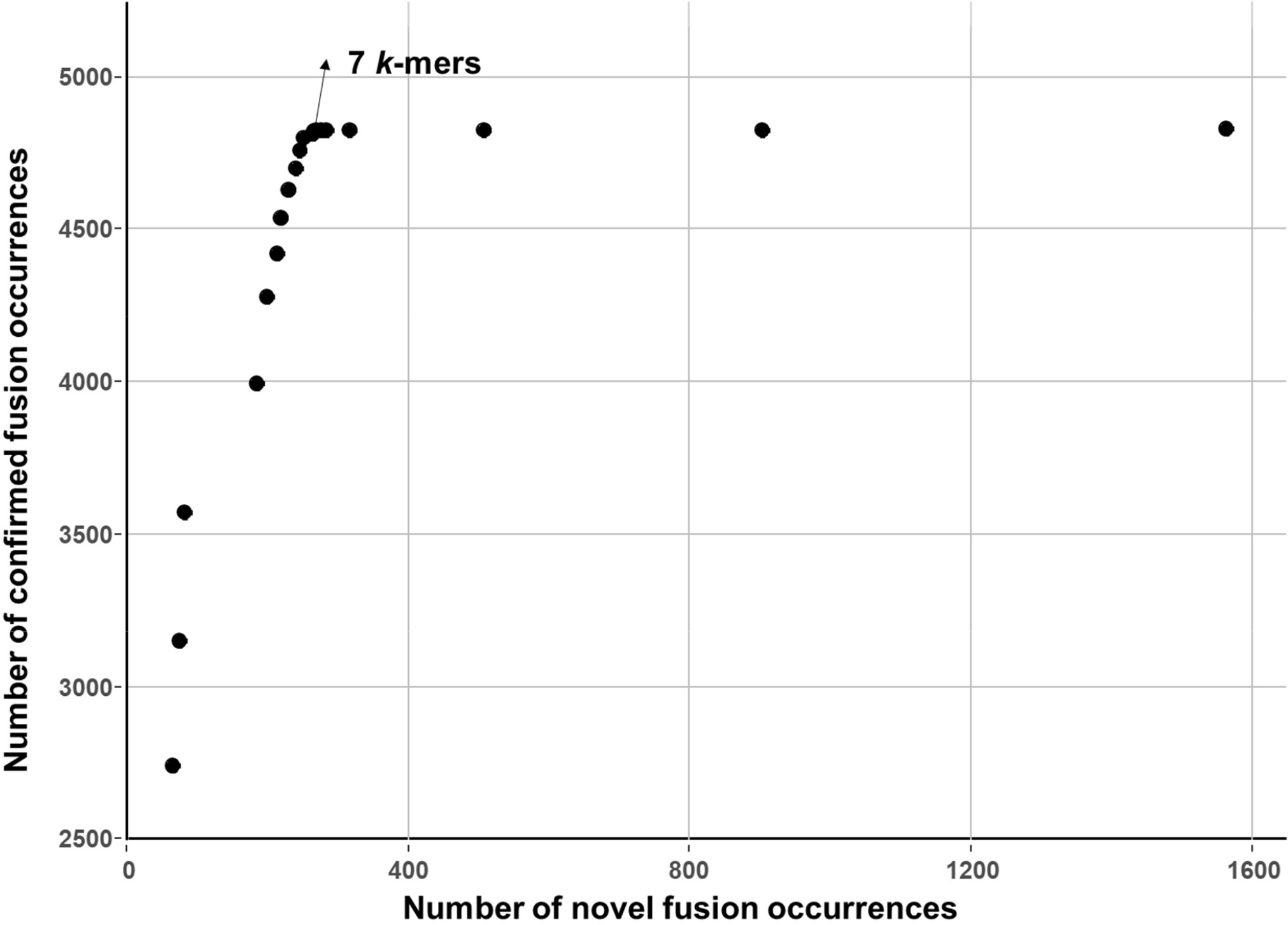
The number of confirmed fusions vs. the number of novel fusions when changing the number of *k*-mer hits required beyond 98^th^ percentile to call a fusion. According to the figure, requiring 7 additional *k*-mers achieved the best balance between sensitivity and specificity and thus it was selected to determine the final set of novel occurrences.

**Supplementary Figure 6.**
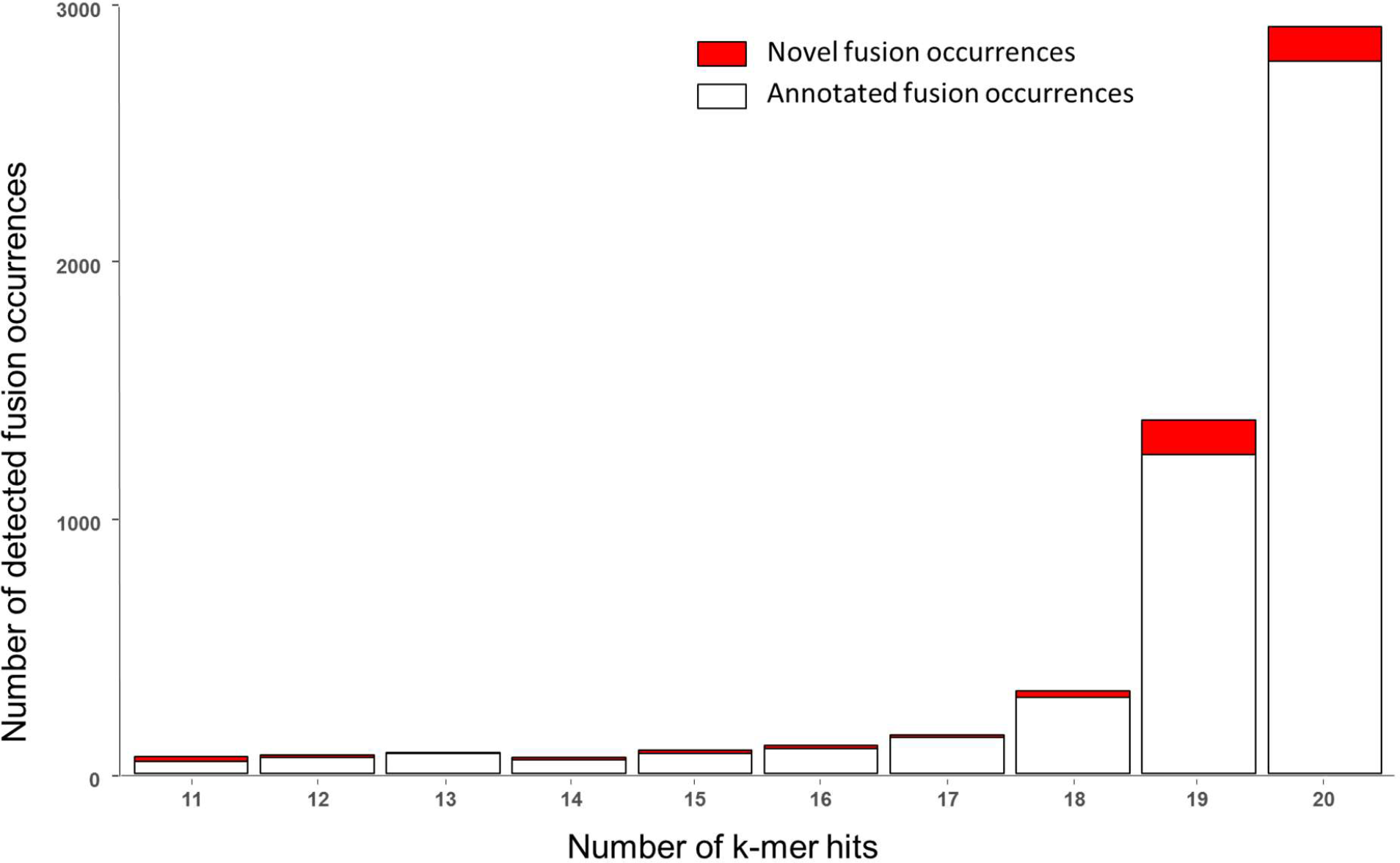
The distribution of the number of *k*-mer hits in annotated and novel fusion occurrences. Most of the fusions carry pretty high *k*-mer hits (18, 19, 20). There do exist quite a number of fusion occurrences in both novel and annotated categories with *k*-mer hits as low as 11 and 12.

**Supplementary Figure 7:**
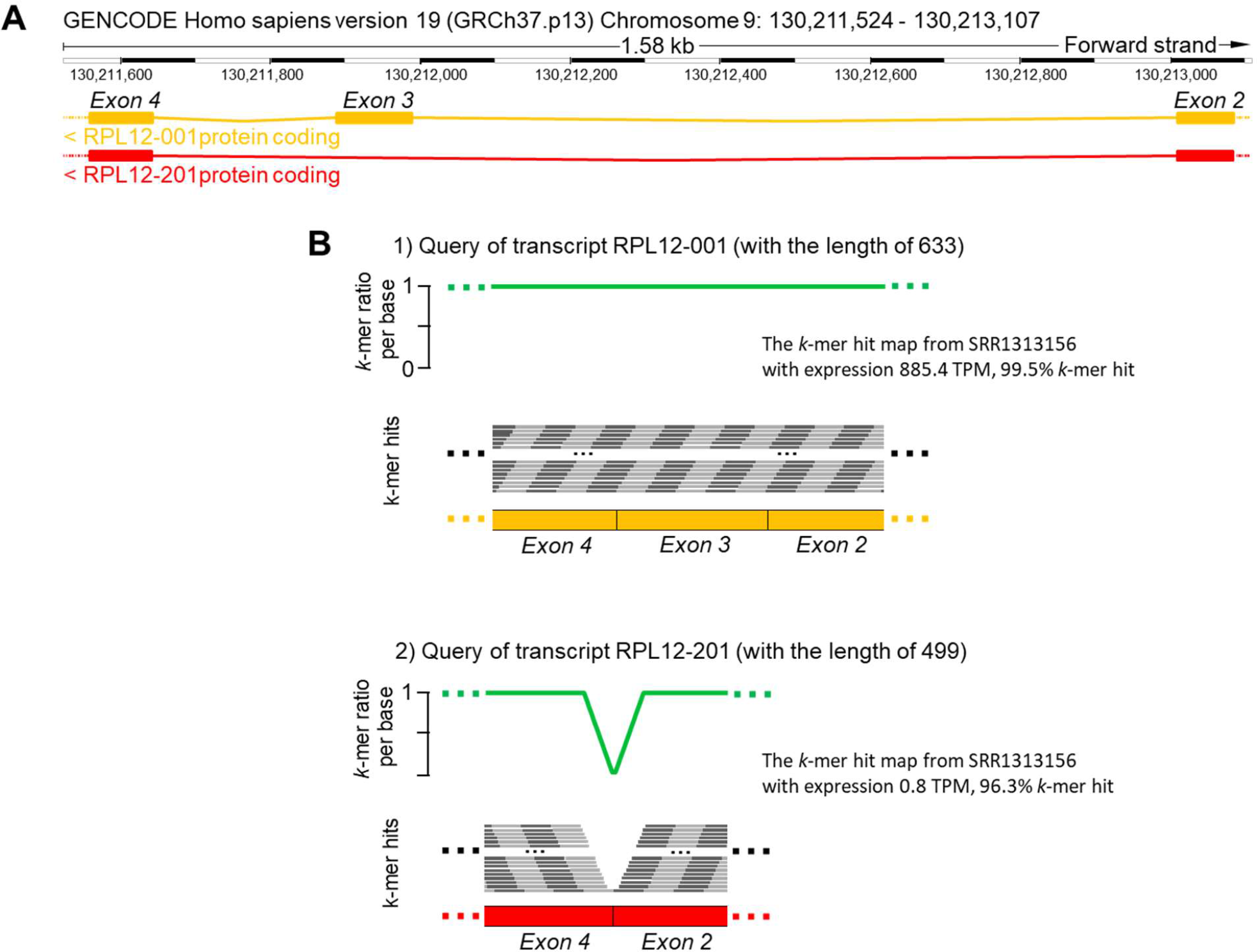
The lack of *k*-mer hits in RNA-seq experiment SRR1313156 connecting exon 2 and exon 4 in transcript RPL12-201 (Shown in B.2) suggests the transcript is barely expressed. This is consistent with its transcript expression estimated by Kallisto using its raw reads. However, the transcript is present with high *k*-mer hit ratio due to the high expression of transcript isoform RPL12-001. As a result, both transcripts will be returned as expressed by SBT-based algorithms should *θ* be set as 0.9.

## Supplementary Tables

**Table 1:** Performance comparison on construction. We compared the time, memory and space usage to construct the SBT^10^, SSBT^11^ and SBT-AS^12^ trees with SeqOthello using the 2,652 SRA RNA-seq samples. The *k*-mer preparation step converts each individual sequencing experiment to the binary format using Jellyfish-based functions with 16 threads. In order to alleviate noise from sequencing errors, *k*-mers having a frequency lower than a specific threshold were removed from the experiment^10^ (see Supp Table 2). The index step follows the k-mer prep step. Unless mentioned otherwise, the comparisons were tested using a single thread. SeqOthello reduces the index construction time by 81% comparing to SBT-AS and the final index size by 32% comparing to the smallest SSBT index.

| Tools | SeqOthello | SBT | SBT-AS | SSBT |
| --- | --- | --- | --- | --- |
| k-mer preparation (**days**) | **3.4** | 4.1 | 4.3 | 4.8 |
| Index building (**hours**) | **1.9** | 39.5 | 10.2 | 46.6 |
| Peak memory (GB) | **14.1** | 23.4 | 39.1 | 5.6 |
| Intermediate disk space (TB) | **0.9** | 1.4 | 3.7 | 1.9 |
| Final index size (GB) | **20.8** | 185.5 | 168.5 | 30.8 |

**Table 2:** The list of k-mer count threshold used to obtain k-mers from Jellyfish as a function of the fasta.gz file size of each experiment. The thresholding criteria are only applied to the 2,652 human RNA-seq experiments from SRA. Only *k*-mers with frequency count no less than the threshold are retained for subsequent indexing.

| fasta.gz size | Threshold |
| --- | --- |
| < 300 MB | 1 |
| between 300 MB and 500 MB | 3 |
| Between 500 MB and 1 GB | 10 |
| Between 1 GB and 3 GB | 20 |
| > 3GB | 50 |

**Table 3:**
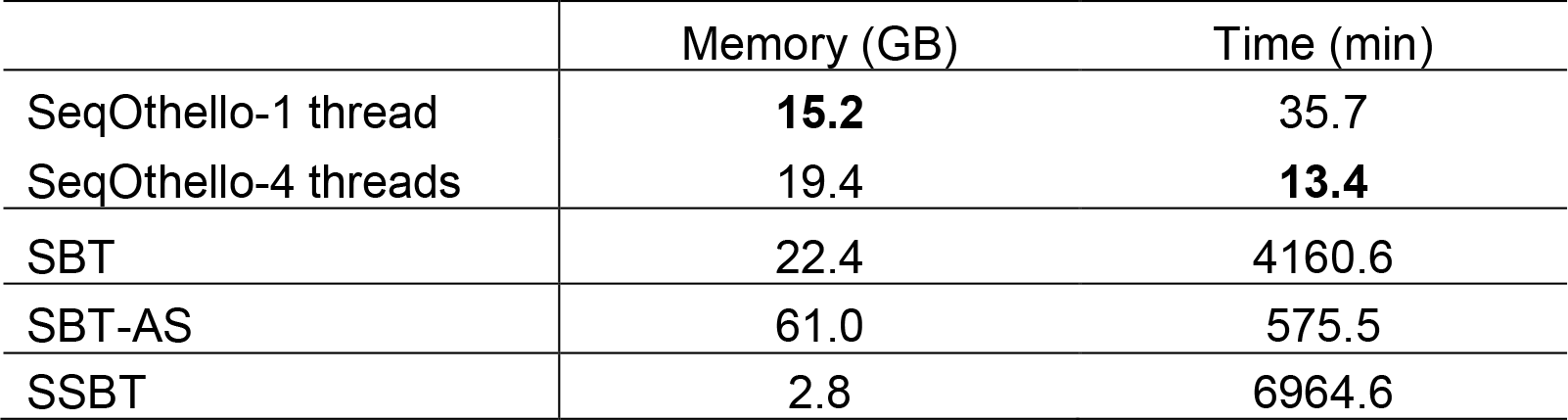
Memory and time usage on large batch query against 2,652 human RNA-seq experiments. We evaluated the performance of SBT, SBT-AS, SSBT and SeqOthello on querying 198,074 transcripts (length >= 20) of the all GenCodeV25 human transcriptome annotations. SBT, SBT-AS and SSBT were tested using Ɵ=0.9 and max-filters = 1.

**Table 4:**
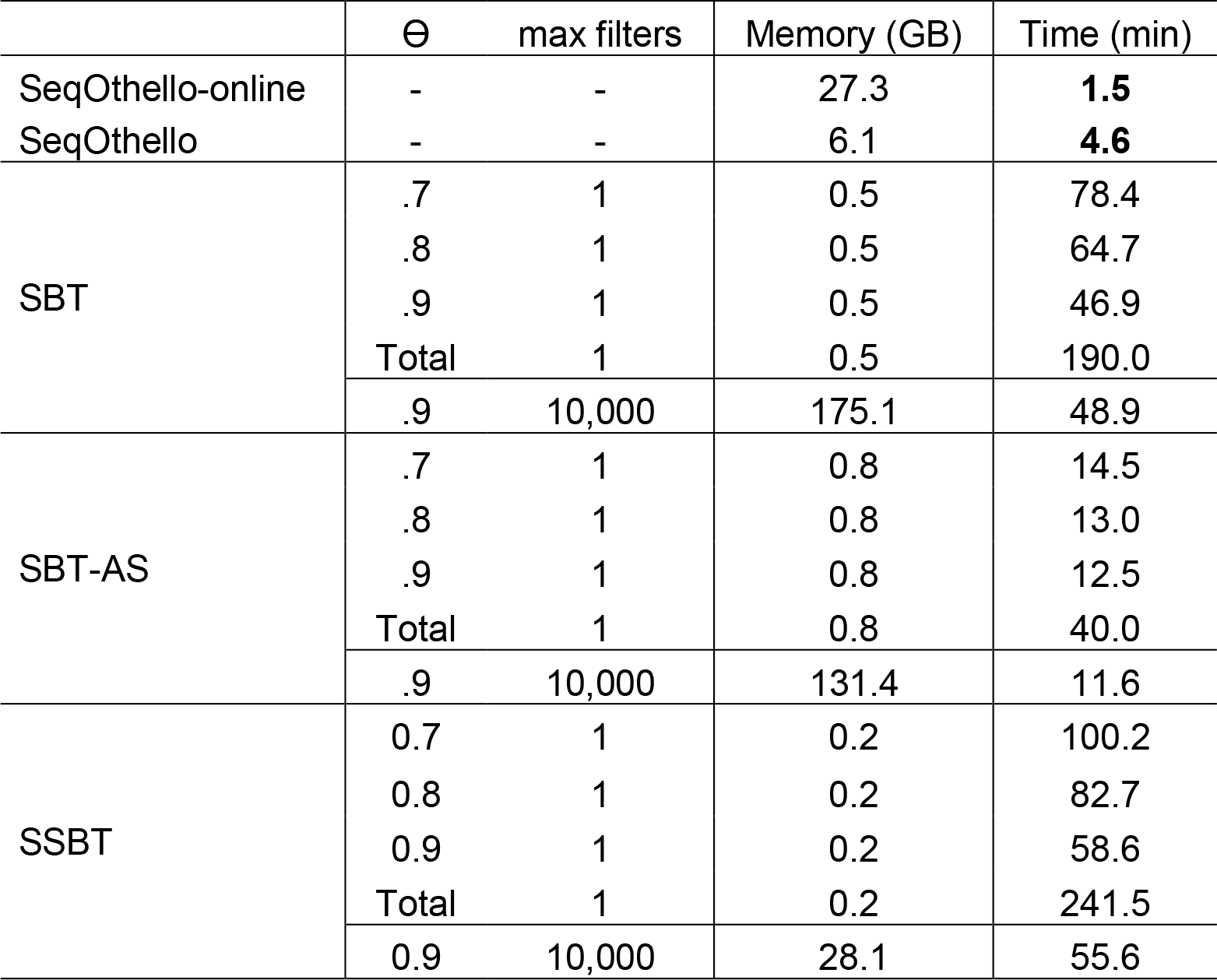
Performance comparison on small batch query. We evaluated the memory and time cost for SBT, SBT-AS, SSBT and SeqOthello on querying 10 sets of 1000 transcripts in the 2,652 experiments. We first estimated the expression profiles of the 198,074 GenCodeV25 transcripts for a randomly selected 200 experiments using Sailfish. We then randomly sampled 10 sets of 1000 transcripts that are expressed above 100 TPM (transcripts per million) in at least one of the experiments to build the small batch query sets. This is to make sure the query sets are non-trival. We benchmarked the query performance of SBT, SBT-AS and SSBT using Ɵ = 0.7, 0.8 and 0.9 with the max-filters setting to 1 and Ɵ = 0.9 with the max-filters setting to 10,000. The max-filters parameter defines the maximum number of filters loaded in the memory for all SBT-based algorithms. Setting this parameter to 10,000 will make sure the entire bloom filter tree staying in the memory without unnecessary eviction. SeqOthello does not require any parameter for sequence query.

